# Generative design of intrinsically disordered proteins based on conditioned protein language models: Data is the limit

**DOI:** 10.64898/2026.04.14.718363

**Authors:** Laure Carrière, Alexandre Huyghe, Mátyás Pajkos, Pau Bernadó, Juan Cortés

## Abstract

Intrinsically disordered proteins and regions (IDRs) are central to a multitude of biological processes. Despite extensive studies of their structural and physicochemical properties, the rational design of IDRs with defined conformational behavior remains challenging due to their ensemble nature. Here we present a generative framework for designing disordered protein sequences conditioned on target conformational ensemble descriptors using protein language models (pLMs). We formulate IDR design as the task of generating amino acid sequences predicted to realize specified biophysical properties and implement a Transformer encoder–decoder architecture that maps numerical descriptors to protein sequences. By training models on datasets spanning two orders of magnitude in size, we show that accurate control of conformational and physicochemical properties is achieved only at large data scale. These results demonstrate the feasibility of conditioning generative models on ensemble-level descriptors for IDR design. More broadly, these results support a data-centric paradigm for protein engineering, in which data availability emerges as a key limiting factor for the accurate design of IDRs.

## Introduction

Computational protein design has experienced rapid progress in recent years, driven by advances in deep learning. These approaches have enabled the generation of novel folded proteins with predefined structures and functions, including enzymes, binders, and self-assembling architectures with high experimental success rates [1–3]. Despite these advances, most efforts have focused on proteins that adopt well-defined three-dimensional structures. In contrast, the rational design of intrinsically disordered proteins and regions (IDRs), which populate heterogeneous conformational ensembles rather than single native states, remains comparatively underdeveloped [4, 5].

IDRs are widespread across proteomes and play essential roles in cellular regulation, signaling, and biomolecular condensation [6–8]. Their functional behavior is often encoded not in a unique folded structure but in ensemble-level properties such as chain compactness, secondary structure propensity, long-range intra-molecular contacts and phase separation propensity. Consequently, design strategies developed for folded proteins cannot be directly transferred to disordered systems. Early approaches to IDR engineering have relied largely on empirical heuristics, including modulation of charge patterning, aromatic content, and hydropathy, which correlate with global conformational properties, such as the radius of gyration or chain compaction [9–11]. While useful, these rules provide limited quantitative control and generality across sequence space.

More sophisticated strategies have emerged based on molecular simulations combined with iterative sequence optimization [12, 13], sometimes coupled with active learning to improve exploration efficiency [14]. These physics-based approaches can capture complex sequence–ensemble relationships but remain computationally expensive, restricting exploration to a tiny fraction of the astronomically large sequence space accessible to disordered proteins. This limitation motivates the development of data-driven generative approaches capable of learning sequence–property relationships directly from large datasets.

We formulate the design of IDRs as the task of generating amino acid sequences that satisfy predefined ensemble-level structural properties, in conceptual analogy to structure-based protein design for folded proteins [3, 15]. In this formulation, sequences are generated to reproduce target descriptors of the conformational ensemble, such as the radius of gyration (*R*_*g*_) or the end-to-end distance (*R*_*ee*_). This framework naturally accommodates additional sequence-based constraints (e.g., net charge, hydrophobicity, or compositional biases) and can be extended to richer representations of conformational behavior, such as residue–residue contact probabilities.

To address this problem, we explored several neural network architectures for conditional sequence generation. The best performance was obtained using a Transformer encoder–decoder framework inspired by the T5 architecture (Text-To-Text Transfer Transformer) [16], in which the encoder processes numerical descriptors of conformational and physicochemical properties, while the decoder autoregressively generates amino acid sequences. Importantly, the encoder and decoder operate on different data representations (i.e. continuous descriptor vectors versus discrete sequence tokens) allowing flexible conditioning on heterogeneous inputs. This design is conceptually analogous to multimodal generative models developed in other domains, such as systems that generate text from images or speech, where distinct encoder and decoder representations are coupled through cross-attention [17]. Our approach is also related to recent protein generative models that condition sequence generation on functional or contextual information, including protein language models (pLMs), such as ProGen [18] and ProtGPT2 [19]. However, those methods typically rely on decoder-only conditioning through discrete tokens, whereas our encoder–decoder formulation enables direct conditioning on continuous biophysical descriptors and provides greater control over the generated sequences.

A central hypothesis of this work is that, as in many areas of deep learning, the performance of generative models for IDR design is strongly constrained by data availability. While large structural databases have been established for folded proteins, most notably the Protein Data Bank (PDB) [20], datasets linking IDR sequences to quantitative descriptors of their conformational ensembles remain extremely scarce. Resources such as DisProt [21] and the Protein Ensemble Database (PED) [22] provide valuable curated annotations, but only cover a limited number of proteins. As a result, recent efforts have begun to rely on computational pipelines to generate large-scale sequence–ensemble annotations [23, 24].

To evaluate the impact of data scale, we trained generative models on two computationally derived datasets differing by approximately two orders of magnitude in size. We show that the proposed framework can generate IDR sequences whose predicted conformational and physicochemical properties closely match the target descriptors provided as input. However, this level of generative control is achieved only when models are trained on large datasets: models trained on more limited data remain capable of producing broadly consistent sequences, but with noticeably reduced accuracy. Together, these results demonstrate both the feasibility of conditioning protein language models on ensemble-level properties for IDR design and the central role of dataset scale in enabling reliable generative performance. More broadly, our findings suggest that large, systematically annotated datasets of disordered proteins and their associated properties will be essential for the next generation of data-driven biomolecular design methods.

## Results

### A Conditioned pLM for IDR Design: IDR-Prop2Seq

The generative framework adopted in this work follows an encoder–decoder architecture in which numerical descriptors encoding conformational and physicochemical properties serve as conditioning inputs. The encoder transforms these descriptors into contextualized representations, while the decoder autoregressively generates amino acid sequences under cross-attention to the encoded signal. The model is implemented as a Transformer architecture inspired by the T5 design [16] and is illustrated in Fig. 1. Conditioning inputs consist of a vector of descriptors associated with each sequence. Instead of concatenating these values into a single representation, each descriptor is projected into a learned embedding and treated as an individual token. This representation allows the encoder to model relationships among descriptors through self-attention. The decoder operates at the amino acid level and generates sequences autoregressively. To enable flexible conditioning, the model also includes learned embeddings representing missing descriptors, allowing sequence generation from partial sets of constraints.

**Figure 1:**
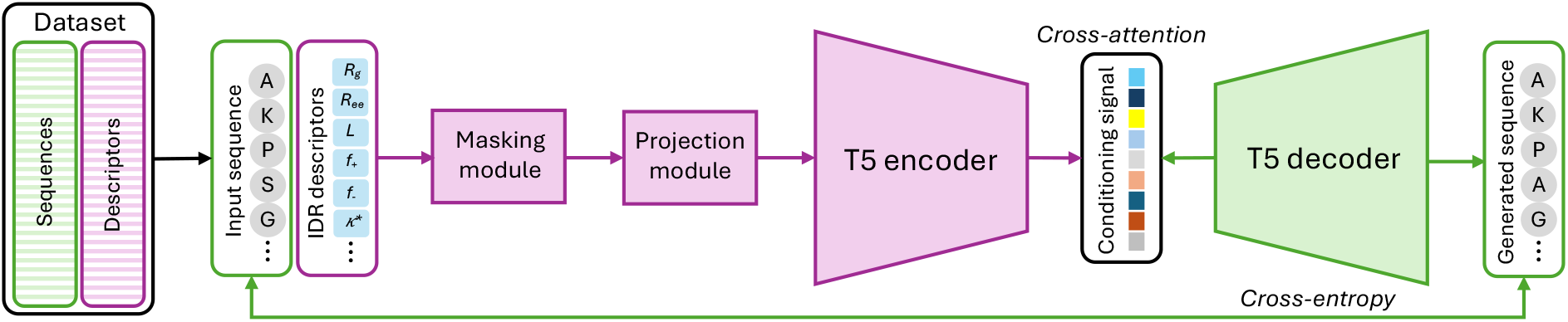
A schematic overview of the architecture and conditioning strategy of the IDP-Prop2seq model. The encoder processes IDR descriptors that define the conditioning signal, while the decoder generates amino acid sequences. Conditioning is achieved through cross-attention between encoder and decoder representations. During training, sequence generation is optimized using a cross-entropy loss.

In this study, we used a vector of 15 descriptors encompassing both conformational and sequence-derived physicochemical properties (see Materials and Methods). This reduced set was chosen to capture a diverse yet practically meaningful subset of IDR characteristics from a user-oriented design perspective. The framework itself is not limited to these descriptors and can readily incorporate additional properties derived from sequence, experiments, or computational predictions.

Models were trained at multiple capacity scales using the same architectural principles and training objective, with hyperparameters adjusted according to the size of the available training data in order to balance model capacity and dataset scale. Training was performed from scratch using standard autoregressive cross-entropy loss with teacher forcing. To improve robustness and allow generation from incomplete inputs, conditioning descriptors were stochastically masked during training, exposing the model to varying combinations of constraints. Full architectural details, hyperparameters, and training procedures are provided in Materials and Methods.

### Datasets

To investigate the impact of dataset scale on conditional sequence generation, we considered two datasets of IDRs differing substantially in size. The first dataset consists of IDRs from the human proteome previously compiled by Tesei *et al*. [23], containing approximately 20,000 sequences; we refer to this dataset as **h-IDRome**. To obtain a substantially larger training resource, we applied a similar identification protocol to multiple bacterial proteomes, producing a dataset of around ten million sequences, hereafter referred to as **b-IDRome**.

For consistency across datasets, all sequences were annotated using the same computational pipeline. Physicochemical descriptors were computed using `idr.mol.feats` [11], and conformational ensemble properties were estimated using `ALBATROSS` [24]. Although conformational descriptors derived from coarse-grained molecular dynamics (CG-MD) simulations are available for the h-IDRome dataset [23], we used predictor-derived values for both datasets to ensure methodological uniformity during training and evaluation. Because the CG-MD-derived descriptors of the human dataset were included in the training set of the `ALBATROSS` predictive model, high predictive accuracy can be expected for these sequences.

Additional dataset construction details are provided in Materials and Methods. In the following, we will name **h**-IDR-Prop2Seq and **b**-IDR-Prop2Seq the models trained from h-IDRome and b-IDRome, respectively.

### Accurate control of conformational properties strongly depends on dataset scale

To assess the capabilities of our generative models, we first evaluated their ability to generate sequences conditioned on a single conformational descriptor, either *R*_*g*_ or *R*_*ee*_. These descriptors are intuitive, experimentally relevant, and directly interpretable from a protein design perspective, making them suitable benchmarks for conditional generation.

To obtain statistically robust estimates, sequence generation was performed in batches of 100 sequences and repeated 1,000 times using random *R*_*g*_ or *R*_*ee*_ values sampled from the empirical distributions observed in the training set. The generated sequences were re-annotated using the same prediction tool. Model performance was quantified by comparing the predicted properties of generated sequences with the corresponding target input values. Fig. 2 shows the distributions of absolute errors for both the `h-IDR-Prop2Seq` and `b-IDR-Prop2Seq` models. A striking difference is observed between the two models: the model trained on the smaller dataset (`h-IDR-Prop2Seq`) exhibits large deviations from the target values, whereas the model trained on the substantially larger dataset (`b-IDR-Prop2Seq`) achieves much smaller errors and tighter distributions. Although both models display outliers, these are more pronounced for the model trained on limited data. Additional analyses indicate that most outliers arise from regions of the descriptor space that are underrepresented in the training data, particularly extreme values (Fig. S1). Together, these results highlight the critical importance of dataset size and coverage for accurate conditional generation of IDR sequences.

**Figure 2:**
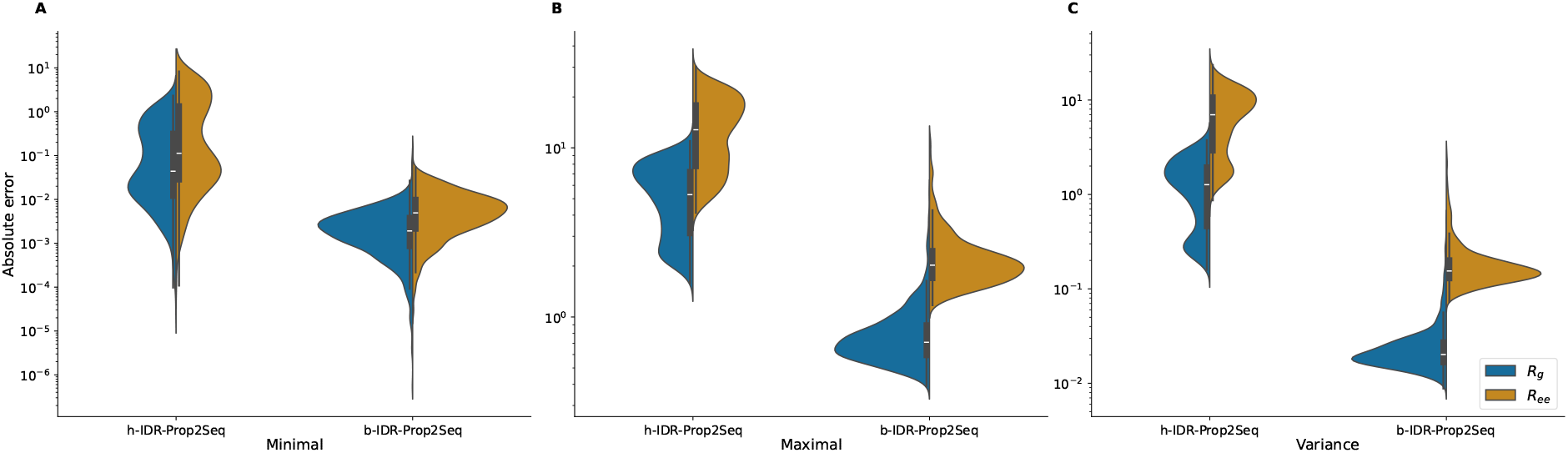
Distribution of absolute error statistics for *R*_*g*_ and *R*_*ee*_. Violin plots showing the distributions of the minimal (**A**), maximal (**B**), and variance (**C**) of the absolute error obtained when generating sequences targeting the conformational descriptors *R*_*g*_ (blue) and *R*_*ee*_ (orange). Results are shown for the `h-IDR-Prop2Seq` model and the `b-IDR-Prop2Seq` model. The vertical axis represents absolute error on a logarithmic scale. For the `b-IDR-Prop2Seq` model, minimal errors are typically in the range of 10^−3^–10^−2^ for *R*_*g*_ and around 10^−2^ for *R*_*ee*_, whereas the `h-IDR-Prop2Seq` model exhibits broader distributions extending from approximately 10^−2^ up to values near 10. Similar trends are observed for maximal errors and variances, which remain substantially lower and more tightly distributed for the model trained on the larger dataset. Boxes indicate the interquartile range with the median shown as a horizontal line.

### Sequence generation remains robust under partial descriptor conditioning

To evaluate the ability of the model to generate sequences when sequence-derived physicochemical constraints are provided in addition to conformational ones, we performed a systematic analysis under partial conditioning. Conditioning vectors were constructed by enforcing the presence of one core descriptor (*R*_*g*_, *R*_*ee*_, or sequence length *L*), together with a random selection of 40% of the remaining descriptors, including both conformational parameters (e.g., *ν, A, R*_0_) and sequence-derived features such as charge content and hydropathy-related descriptors (see Materials and Methods for additional details on the considered descriptors). A total of 1,000 such masks were applied to randomly selected IDR descriptor vectors from the test set. Sequences were generated with the `b-IDR-Prop2Seq` model and evaluated by predicting the complete set of descriptors using the same tools employed during dataset construction.

Model performance was quantified using the variance-normalized mean absolute error (NMAE, see Material and Methods), allowing direct comparison across descriptors with different scales. The resulting distribution of errors (Fig. S2) shows that the model generally maintains good control of the target properties even when conditioning information is incomplete. The median NMAE is approximately 0.29, with 75% of the samples below 0.52 and 90% below 0.92. However, the distribution exhibits a long tail, with a small fraction of cases reaching larger errors (up to ∼2.6 at the 99th percentile).

Inspection of the high-error cases reveals two main contributing factors. First, large errors tend to occur for descriptor values that are poorly represented in the training data, consistent with the behavior observed for the *R*_*g*_ and *R*_*ee*_ targets in the previous analysis. Second, error levels depend strongly on which subsets of descriptors are provided as input. In particular, certain combinations of descriptors lead to systematically larger deviations, reflecting the fact that some properties (e.g., conformational parameters such as *R*_0_ and *ν*) are more difficult to satisfy jointly with other constraints, even when the conditioning vectors are sampled from real IDR examples. Additional analyses presented in Supporting Information (Fig. S3) further illustrate this effect by showing that the normalized error varies substantially across descriptor pairs. Together, these results highlight both the robustness of the model under partial conditioning and the importance of carefully selecting descriptor combinations when defining target property vectors for generative design.

### Wide sequence space coverage and high diversity

We next assessed how broadly the model explores sequence space relative to the original training data, and how diverse the sequences are. Analyses are presented for the `b-IDR-Prop2Seq` model using conditioning on a single conformational descriptor, either *R*_*g*_ or the *R*_*ee*_.

Sequence space coverage was visualized by embedding all sequences using the XL-ProtT5 encoder pretrained on UniRef50 [25], followed by two-dimensional projection with PacMap [26] (see Materials and Methods). The resulting maps (Fig. S4) show that generated sequences populate regions that largely overlap with the density of training sequences. This indicates that the model does not remain confined to a limited subset of sequence space but instead explores a broad portion of the manifold defined by the training data.

Sequence diversity was quantified using SHARK [27], an alignment-free similarity metric well suited for disordered or difficult-to-align sequences (see Materials and Methods). Fig. 3 reports the distributions of maximal and median SHARK similarity scores computed (*i*) within each batch of 100 generated sequences and (*ii*) between generated sequences and training-set sequences with matching *R*_*ee*_ values within a narrow tolerance window chosen to retrieve approximately 100 reference sequences. Across the 1,000 generations conditioned on *R*_*ee*_, similarity distributions are centered near zero, indicating high sequence variability both within batches and relative to the training set (the equivalent analysis for *R*_*g*_ is shown in Fig. S5). Although a small fraction of sequences exhibit moderate similarity either within the same batch or with training-set sequences, the large majority share less than 40% similarity.

**Figure 3:**
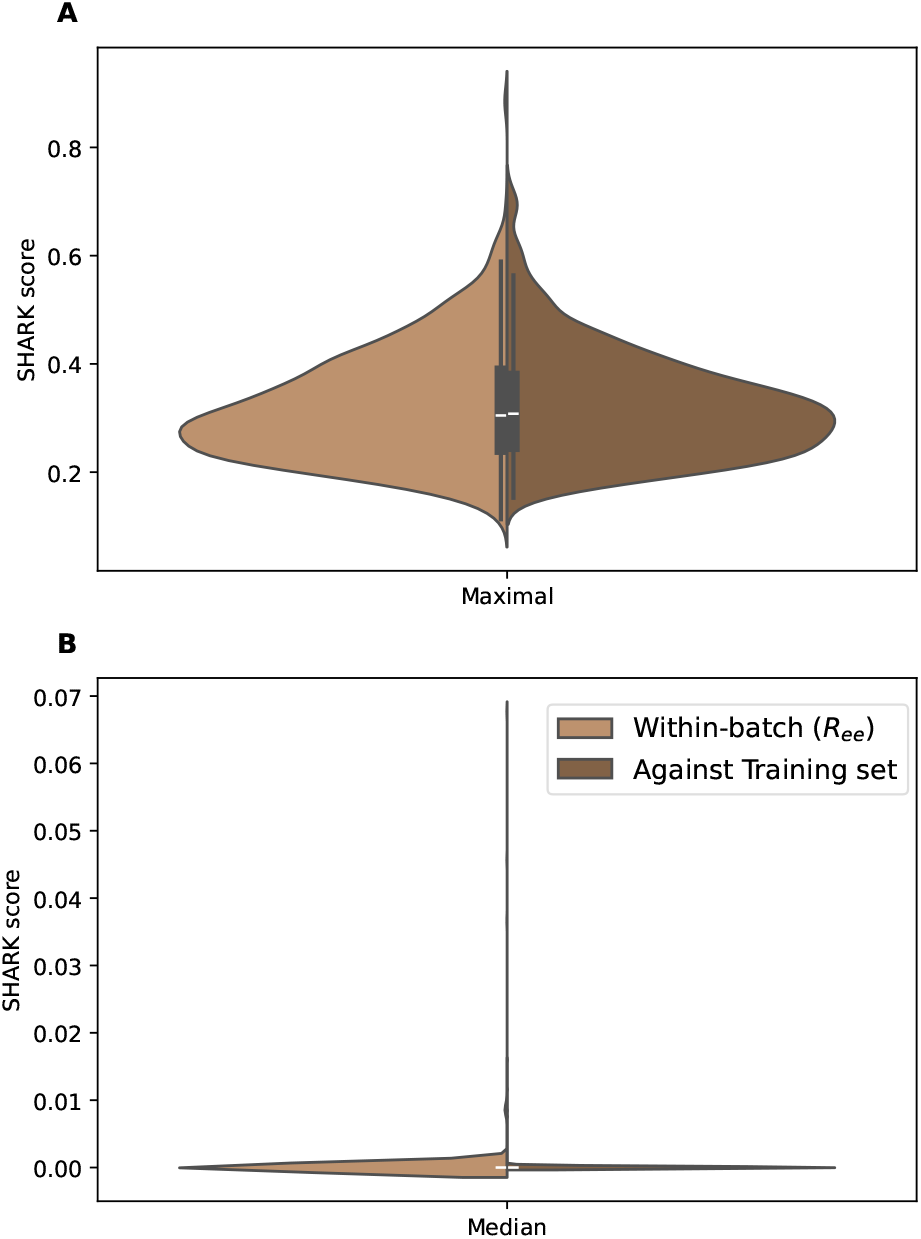
Distribution of SHARK similarity scores for sequences generated in batches of 100 sequences across 1,000 generations conditioned on *R*_*ee*_. Panels show the distributions of the (**A**) maximal and (**B**) median SHARK scores. Light brown violins report similarities computed within each generation batch, whereas dark brown violins report similarities between generated sequences and training-set sequences with matching *R*_*ee*_ values. Overall, similarity values are low, indicating high diversity among generated sequences and no overlap with training sequences.

Overall, these analyses show that generated sequences remain broadly consistent with the sequence space defined by the training data while maintaining substantial diversity both within batches and relative to existing sequences. Note that the balance between sequence diversity and proximity to the training distribution can be further modulated at inference time through decoding parameters such as temperature or sampling thresholds. These results support the use of the model for exploratory design, where both coverage of diverse sequence candidates and avoidance of redundant outputs are important.

## Discussion

This work provides a proof of concept that conditioned pLMs can be used to generate IDR sequences that satisfy user-defined conformational and physicochemical descriptors. By combining a Transformer encoder–decoder architecture with ensemble-level conditioning, we demonstrate that sequence generation can be guided toward targeting conformational properties, extending recent advances in data-driven protein design to disordered systems.

A central finding of this study is that dataset size is a primary determinant of model performance. Training on only a few thousand sequences was insufficient to achieve accurate control of target descriptors, whereas models trained on datasets larger by approximately two orders of magnitude produced sequences that closely matched the desired properties. This observation highlights the importance of data scale for learning sequence–ensemble relationships and suggests that limitations in currently available annotated datasets represent a major bottleneck for the development of reliable generative models for IDRs. Systematic evaluation across intermediate dataset sizes would provide valuable insights into scaling behavior and data efficiency, although such analyses require substantial computational resources.

More broadly, our results support a data-centric perspective in which expanding and improving datasets may yield greater gains than increasing architectural complexity alone.

The datasets used in this work rely on predictive methods at multiple levels, including identification of disordered regions from sequence and estimation of conformational descriptors. Although disorder predictors have improved substantially in recent community-wide assessments [28] and ensemble prediction methods are rapidly advancing [29, 30], uncertainties remain, particularly for quantitative conformational properties derived from sequence alone [31, 32]. Errors introduced at either stage may propagate into the training data and ultimately affect model performance.

In the present study, we deliberately focused on relatively simple one-dimensional descriptors of conformational ensembles: *R*_*g*_ and *R*_*ee*_. These global observables provide intuitive and experimentally relevant measures of chain expansion and compaction, making them attractive targets for an initial exploration of conditional sequence generation. However, they represent only a coarse description of the underlying conformational ensemble. Achieving finer control over IDR behavior will likely require richer representations, for example descriptors based on residue–residue distance distributions or contact probabilities [33]. Generating such descriptors for large training datasets remains challenging, as it requires predictive modeling approaches that are both sufficiently accurate and validated against biophysical data while remaining scalable to high-throughput applications.

Several simplifying assumptions were made in the present framework. The approach considers isolated disordered regions and does not explicitly account for the influence of neighboring folded domains, intermolecular interactions, environmental conditions, or post-translational modifications, all of which can modulate conformational behavior *in vivo*. Incorporating contextual information represents an important direction for future work. For example, conditioning models on environmental variables such as ionic strength or temperature, or integrating sequence context from multi-domain proteins, could enable more realistic and application-specific designs. Similarly, extending the descriptor space to include interaction and phase separation propensities would broaden the range of accessible functional properties of the designed sequences.

Despite these limitations, the proposed generative framework may already be useful for practical applications. One immediate opportunity is the design of disordered linkers connecting protein domains in synthetic and biotechnological constructs, where properties such as flexibility, compaction, and effective inter-domain spacing are critical determinants of function.

In conclusion, this study demonstrates that conditioning pLMs on ensemble-level descriptors enables controllable generation of IDR sequences and identifies data availability as a key limiting factor for progress. As larger and more accurate datasets linking sequence to conformational behavior become available, generative approaches are likely to play an increasingly important role in the rational engineering of disordered proteins and their functions.

## Materials and Methods

### Datasets

The **h-IDRome** dataset was derived from the repository reported by Tesei *et al*. [23] (updated December 2023). Sequences longer than 150 residues were excluded, yielding 20,329 entries.

To construct a large-scale collection of bacterial IDRs (**b-IDRome**), 9,097 bacterial reference proteomes were obtained from UniProt. Redundancy reduction was performed at both the protein and region levels to limit the overrepresentation of homologous proteins and highly similar IDRs across the large number of bacterial proteomes analyzed. All protein sequences were pooled and clustered at the protein level using MMseqs2 [34] with a sequence identity threshold of 70% and minimum coverage of 80%, retaining one representative sequence per cluster. The resulting non-redundant protein set was used as the starting point for disorder annotation. AlphaFold [35] structural predictions for the representative proteins were retrieved from the AlphaFold Protein Structure Database [36], and IDRs were identified using AlphaFold confidence-based disorder annotation, analogous to the **h-IDRome** approach. More precisely, residues were classified according to their per-residue AlphaFold confidence score (pLDDT): residues with pLDDT *>* 80 were labeled as well-folded, residues with pLDDT *<* 70 as disordered, and residues with 70 ≤ pLDDT ≤ 80 as gap regions. To reduce spurious short segments, well-folded and disordered regions shorter than 10 residues were reclassified as gaps. Gap regions were subsequently reassigned based on their sequence context, such that gaps flanked by disordered regions were relabelled as disordered, whereas remaining gaps were considered well-folded. Continuous disordered segments of at least 10 residues were extracted as candidate IDRs. To construct a non-redundant dataset, the extracted IDRs were further redundancy-reduced at the sequence level using MMseqs2 clustering at 90% sequence identity and 90% coverage, retaining one representative sequence per cluster. As a final step, to mitigate length bias caused by the over-representation of short IDRs (10–20 residues), half of these regions were randomly removed, while regions longer than 128 residues were excluded. The resulting dataset constitutes a non-redundant bacterial IDRome comprising 10,821,730 IDRs.

### IDR Descriptors

The descriptors used in this study fall into two categories: conformational descriptors, which characterize properties of the conformational ensemble, and sequence-derived descriptors, which capture physicochemical and compositional features that can be computed directly from the amino acid sequence. Conformational descriptors were obtained from `ALBATROSS` [24] and include five ensemble-averaged quantities: the radius of gyration (*R*_*g*_), the end-to-end distance (*R*_*ee*_), the Flory scaling exponent (*ν*), the asphericity (*A*), and the scaling prefactor (*R*_0_). These parameters provide complementary information about chain expansion, shape, and polymer behavior. Sequence-derived descriptors, computed using idr.mol.feats [11], were selected based on their known influence on conformational compaction and expansion. These include the sequence length (*L*); charge-related metrics such as the fraction of charged residues (FCR), the net charge per residue (NCPR), the fractions of positively (*f*_+_) and negatively (*f*_−_) charged residues, and charge patterning descriptors including sequence charge decoration (SCD), *κ*∗, and *ω*∗; as well as hydropathy-related features, including sequence hydropathy decoration (SHD) and the Kyte-Doolittle mean hydropathy (⟨*H*⟩).

Together, these quantities define a descriptor vector 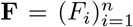 with *n* = 15, which serves as the conditioning input for the generative model.

### Model Architecture

We explored several neural network architectures for conditional sequence generation (details not shown). Preliminary experiments led us to adopt a Transformer-based encoder–decoder architecture [17] inspired by T5 [16]. The model consists of stacked self-attention and feed-forward blocks with residual connections and layer normalization. In this framework, the encoder processes the conditioning descriptors through multi-head self-attention layers to produce contextualized representations. The decoder generates amino acid sequences autoregressively using masked self-attention (causal masking), encoder–decoder crossattention, and position-wise feed-forward networks. Unlike the original T5 formulation, which operates on discrete text tokens, our encoder receives continuous embeddings derived from numerical descriptors.

### Conditioning Representation and Sequence Decoding

Each training sequence is associated with a descriptor vector **F**. Rather than concatenating all descriptors into a single embedding, the conditioning signal is represented as a sequence of length *n*, where each descriptor is mapped to an individual embedding token. For each descriptor value *F*_*i*_, a learned projection Linear(1 → *d*_model_) is applied, followed by a GELU non-linearity, dropout, and layer normalization. The resulting embeddings are provided to the encoder, which models relationships between descriptors through self-attention.

To enable partial conditioning, we introduce a learned descriptor-specific embedding used whenever a value is missing. If a descriptor is present, its projected value is used; otherwise it is replaced by the corresponding learned “missing-descriptor” embedding. This mechanism explicitly distinguishes between a descriptor value equal to zero and an unspecified descriptor.

The decoder generates amino acid sequences autoregressively from the conditioning representations through encoder-decoder cross-attention. Sequences are tokenized at the amino acid level (20 canonical residues) with three special tokens: pad, eos, and unk, yielding a vocabulary of 23 tokens.

### Training Procedure

The datasets were split into training, validation, and test subsets. We used an 85/10/5 split for h-IDRome and an 87/10/3 split for b-IDRome, slightly increasing the proportion of training data relative to the commonly used 80/10/10 partition in order to favor model learning.

To train models capable of generating sequences from incomplete constraints, stochastic masking was applied to conditioning descriptors during training. The primary conformational descriptors, *R*_*g*_ and *R*_*ee*_, and sequence length *L* were treated as core variables. In most cases, exactly one of these three descriptors was retained and the other two masked; with probability 0.15, two were retained instead. Remaining descriptors were independently masked with probability 0.6. Masked descriptor were set to zero in normalized space and flagged as absent via a *mask_presence* tensor, triggering the use of the learned missing-descriptor embedding.

Models were trained using autoregressive cross-entropy loss between predicted token distributions and ground-truth amino acid tokens under teacher forcing. Padding positions were excluded from the loss by setting corresponding labels to −100. Label smoothing was set to 0.

Optimization was performed using fused AdamW (*adamw_torch_fused*) with learning rate 10^−4^ and weight decay 0.01. A cosine learning-rate schedule with restarts (*cosine_with_restarts*) was used with 2 cycles and a warmup ratio of 0.10. Early stopping based on validation loss was applied with a patience of 6 epochs, and the best checkpoint was retained.

### Model Capacity and Computational Setup

During model development, several architectural configurations and conditioning strategies were explored, including alternative tokenization schemes, different mechanisms for incorporating numerical descriptors, and multiple Transformer capacities. Model variants were first evaluated based on validation loss during autoregressive training and subsequently assessed based on conditional generation performance, measured by the agreement between target descriptors and the predicted properties of generated sequences.

Based on these preliminary experiments, we retained two model variants differing only in Transformer capacity while keeping the conditioning mechanism and decoding objective identical. Both models use 8 encoder and 8 decoder layers. The `h-IDR-Prop2Seq` model uses a hidden dimension of 512 with 4 attention heads and a feed-forward dimension of 1024, while the larger `b-IDR-Prop2Seq` model uses a hidden dimension of 1024 with 16 attention heads and a feed-forward dimension of 3072. This choice allowed us to balance model expressiveness with dataset size while maintaining comparable training procedures across experiments. These configurations correspond to 29.4 million trainable parameters for `h-IDR-Prop2Seq` and 201.4 million for `b-IDR-Prop2Seq`.

All training runs were performed on a single NVIDIA H100 GPU using mixed precision. Gradient accumulation was employed to achieve large effective batch sizes: 1024 for the `h-IDR-Prop2Seq` model and 4096 for the `b-IDR-Prop2Seq model`.

### Generation and Evaluation under Descriptor Conditioning

Sequences were generated by conditioning the model on numerical descriptors defining conformational or physicochemical properties. At inference time, generation was performed using stochastic decoding with a temperature of 1.0 and nucleus sampling (*top-p*=0.9), without *top-k* filtering. A repetition penalty of 1.1 was applied to discourage trivial repetition. Two conditioning scenarios were considered.

In the first case, generation was performed using a single conformational descriptor, either *R*_*g*_ or *R*_*ee*_. For each dataset, empirical distributions of these descriptors were estimated from the training set, and 1,000 values were sampled to construct conditioning inputs. For each conditioning value, 100 sequences were generated with each trained model.

In the second case, generation was performed under partial conditioning using multiple descriptors. A total of 1,000 partially masked descriptor vectors were constructed from the test set, each including one mandatory core descriptor (*R*_*g*_, *R*_*ee*_, or sequence length *L*) combined with a random selection of 40% of the remaining descriptors.

In all cases, generated sequences were evaluated by re-estimating their descriptors using the same predictive tools employed during dataset construction (`ALBATROSS` and `idr.mol.feats`). Performance was quantified using the variance-normalized metric described below, enabling comparison across heterogeneous descriptor sets.

### Normalized Mean Absolute Error (NMAE)

To quantify the agreement between requested and generated properties, we defined a normalized mean absolute error (NMAE). Let **F**^ref^ denote the vector of descriptor values used as conditioning input and **F**^gen^ the corresponding vector predicted for the generated sequence. For each descriptor *F*_*i*_, the normalized error is defined relative to the standard deviation *σ*_*i*_ of that descriptor in the training set:

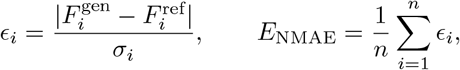

where the sum runs over the *n* unmasked descriptors. Normalization by *σ*_*i*_ makes the metric dimensionless and directly comparable across heterogeneous descriptors. As a result, descriptors with low intrinsic variability in the training set contribute more strongly to the error for a given absolute deviation.

Values of *E*_NMAE_ *<* 1 indicate that generated properties deviate on average by less than one standard deviation from the target descriptors, whereas values approaching 0 correspond to near-perfect agreement.

### Sequence Space Coverage and Diversity

To evaluate how generated sequences populate sequence space relative to the original datasets, we embedded all sequences using the XL-ProtT5 encoder trained on UniRef50, as implemented by Elnaggar *et al*. [25], yielding 1,024-dimensional representations. These embeddings were projected into two dimensions using PacMap [26] to visualize global coverage patterns, with the number of neighbors set to 20, the MN ratio to 0.25, the FP ratio to 10, and the number of iterations to 400.

Sequence-level similarity was quantified using SHARK (Similarity/Homology Assessment by Relating K-mers) [27], a k-mer–based method with performance comparable to local alignment approaches. Sequences were decomposed into overlapping k-mers of length *k* = 7. Pairwise similarity was computed by considering all k-mer pairs exceeding a similarity threshold of 0.9 and aggregating these contributions into a final SHARK score.

## Data and Software availability

All datasets, source code, and trained model weights necessary to reproduce the results of this study will be made publicly available upon publication of the article.

*Acknowledgements*

This work was supported by the French National Research Agency (ANR) under grant ANR-22-CE45-0003 (CORNFLEX project) and by the Occitanie Region as part of the “AI for Health” Program. It was also carried out in the framework of the COST Action ML4NGP [CA21160], supported by the COST (European Cooperation in Science and Technology) and the HORIZON-MSCA-SE project IDPfun2, funded by the European Union under grant agreement no. 101182949.

## Notes

### Competing Interest Statement

The authors have declared no competing interest.

