## Supplementary figures for "Generative design of intrinsically disordered proteins based on conditioned protein language models: Data is the limit"

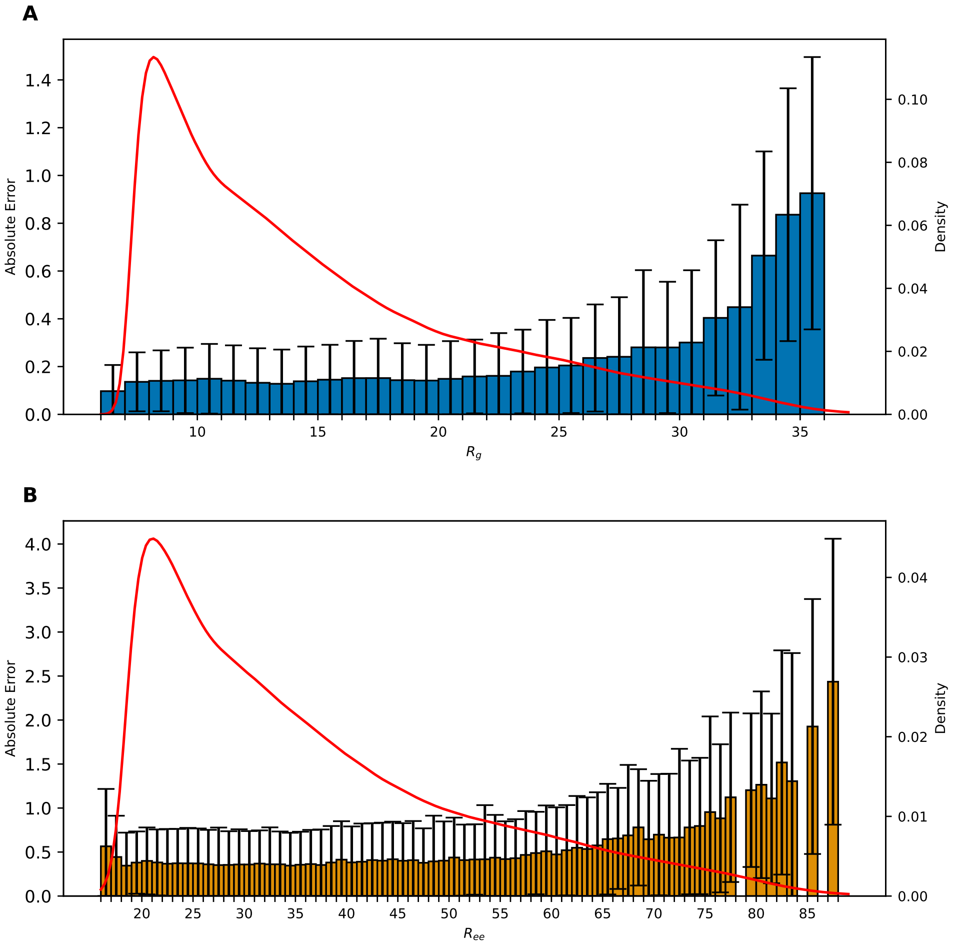


Figure 1: Absolute error distributions for sequences generated with `b-IDR-Prop2Seq` as a function of the conditioning value, using (**A**) $R_g$ or (**B**) $R_{ee}$. For each sampled descriptor value, the distribution of absolute errors across generated sequences is shown, and black error bars indicate the corresponding standard deviation. The red curves report a kernel density estimate of the training-set distribution of $R_g$ or $R_{ee}$, highlighting regions of descriptor space that are well or poorly represented in the training data. In both cases, larger errors are primarily observed for extreme descriptor values, consistent with reduced training-set coverage.

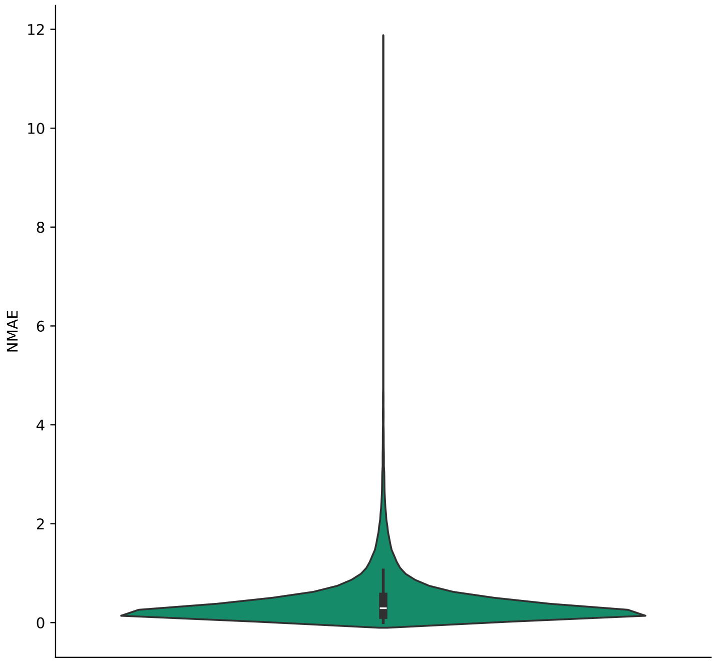


Figure 2: Distribution of the normalized mean absolute error (NMAE) obtained for the `b-IDR-Prop2Seq` model under partial conditioning. NMAE values quantify the deviation between target and generated descriptor vectors, normalized by descriptor variability in the training set (see Methods). The distribution is centered at low values (median $\approx 0.29$), indicating that the model generally preserves the requested properties despite incomplete conditioning. Approximately 91% of the samples have NMAE $< 1$, corresponding to an average deviation of less than one standard deviation across unmasked descriptors.

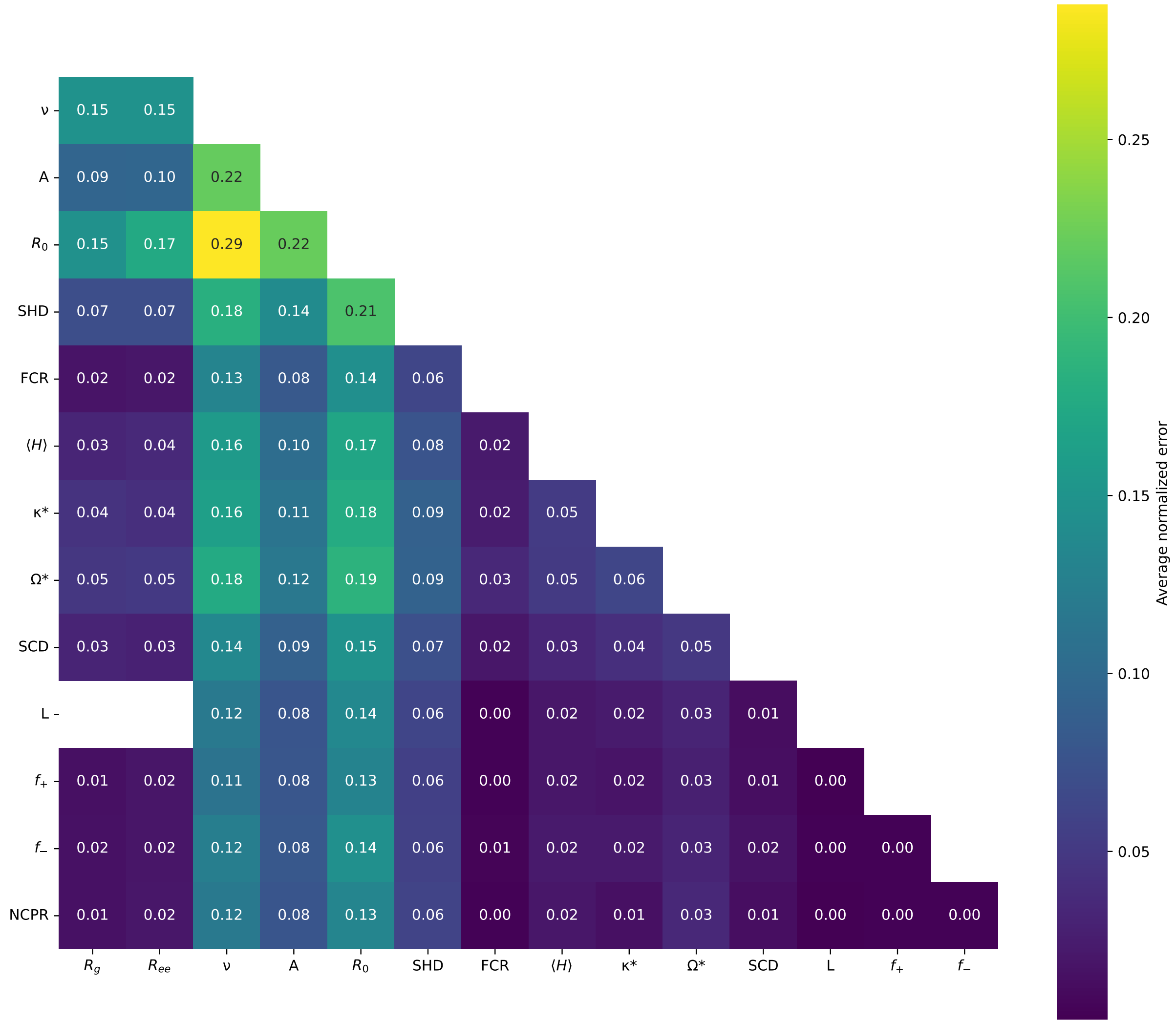


Figure 3: Pairwise analysis of descriptor interactions under partial conditioning for the `b-IDR-Prop2Seq` model. The heatmap reports the average normalized error $\epsilon_i$ (see Methods) for generations in which pairs of descriptors co-occur in the conditioning input. Blank entries correspond to descriptor pairs that never appear together in the masking protocol (e.g., $R_g$, $R_{ee}$, and $L$, which were enforced as mutually exclusive core descriptors). Conditioning vectors including both $R_0$ and $\nu$ tend to yield higher normalized errors, consistent with increased difficulty in simultaneously satisfying these correlated conformational constraints. More generally, partial inputs containing either $R_0$ or $\nu$ are associated with larger $\epsilon_i$ values compared with other descriptor combinations.

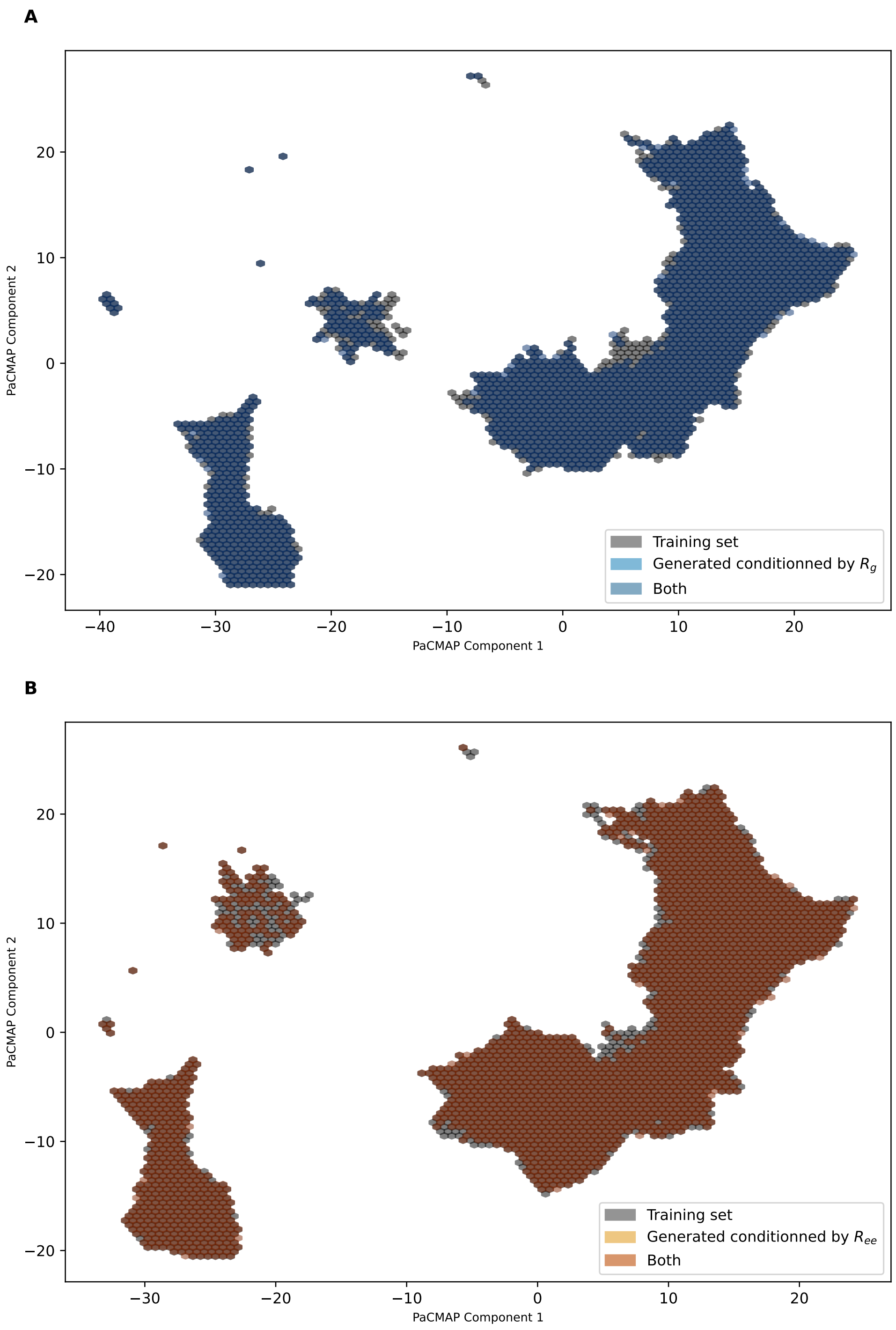


Figure 4: Sequence space coverage of sequences generated by `b-IDR-Prop2Seq` under single-descriptor conditioning. Training sequences (gray) and generated sequences (colored) are embedded using XL-ProtT5 representations [?] and projected into two dimensions with PacMap [?]. Panels show generations conditioned on (**A**) $R_g$ and (**B**) $R_{ee}$. Generated sequences broadly overlap with the training-set manifold, indicating wide coverage of sequence space under both conditioning regimes.

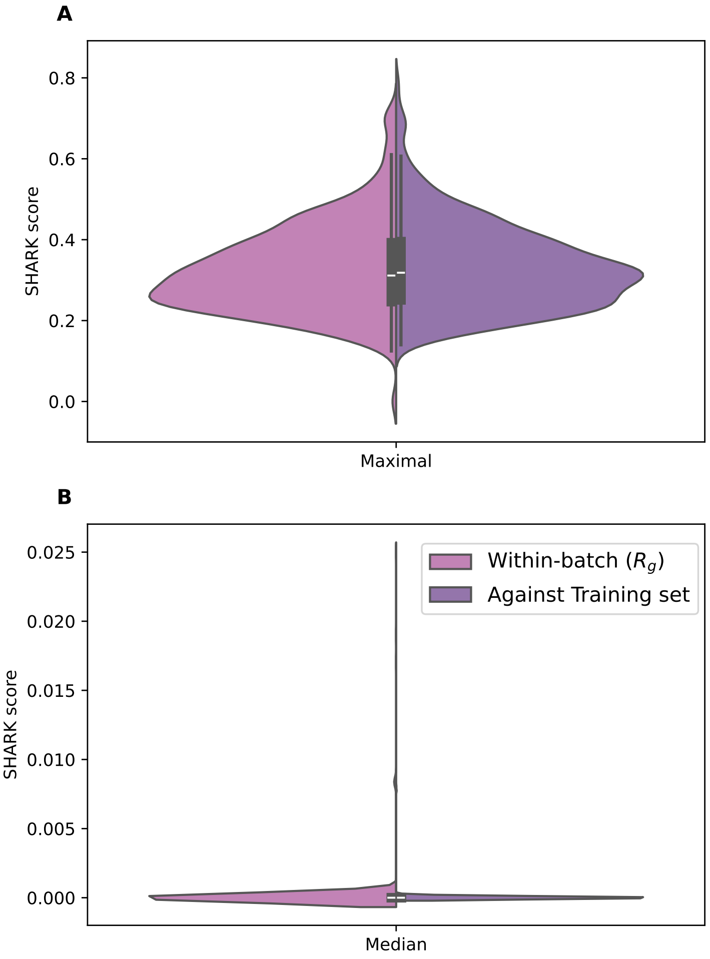


Figure 5: Distribution of SHARK similarity scores for sequences generated in batches of 100 sequences across 1,000 generations conditioned on $R_g$. Panels show the distributions of the (**A**) maximal and (**B**) median SHARK scores. Pink violins report similarities computed within each generation batch, whereas violet violins report similarities between generated sequences and training-set sequences with matching $R_g$ values. Overall, similarity values remain low, indicating high diversity among generated sequences and no overlap with training sequences. In particular, more than 75% of the sequences exhibit less than 40% similarity both within batches and relative to the training set.
